# Exposing the core genetic drivers of chronological aging in yeast cells

**DOI:** 10.1101/2025.03.23.644847

**Authors:** Erika Cruz-Bonilla, Sergio E. Campos, Soledad Funes, Cei Abreu-Goodger, Alexander DeLuna

## Abstract

The chronological lifespan of *Saccharomyces cerevisiae* has been pivotal in advancing our understanding of aging in eukaryotic cells. However, gaining a genome-wide perspective of this trait remains challenging due to substantial discrepancies observed across large-scale gene-deletion screens. In this study, we performed a meta-analysis to compile a ranked catalog of key processes and regulators driving chronological longevity in yeast, ensuring their robustness across diverse experimental setups. These consistent chronological aging factors were enriched in genes associated with yeast replicative lifespan and orthologs implicated in aging across other model organisms. Functional analysis revealed that the downstream cellular mechanisms underlying chronological longevity in yeast align with well-established, universal hallmarks of aging, underscoring the potential of the yeast chronological aging model to investigate conserved aging processes. Additionally, we identified transcriptional regulators associated with these consistent genetic factors, uncovering potential global and local modulators of chronological aging. Among these, Tec1, a key regulator of the filamentous growth pathway, emerged as a central hub connected to multiple aging pathways. To further elucidate the functional role of this regulator, we conducted a high-resolution lifespan-epistasis screen, demonstrating that *TEC1* and mitochondrial machinery promote chronological longevity in parallel, compensating for each other’s impaired functions. Our findings provide an integrated view of the core genetic and functional landscape underlying aging in yeast cells.

## Introduction

Lifespan is a complex biological trait shaped by the rate of aging, which is intricately influenced by both genetic and environmental factors [1]. Model organisms have been instrumental in uncovering the complex genetic mechanisms underlying aging and longevity. Notably, the *age-1* phosphoinositide 3-kinase that regulates the insulin/IGF-1 signaling pathway, was the first “aging gene” identified in *Caenorhabditis.* The sole mutation of this gene extends lifespan by up to 65% [2]. Similarly, mutations in the insulin/IGF-1 receptor gene *daf-2* extend the lifespan of *Drosophila* [3], while mice lacking the insulin receptor in adipose tissue also live longer [4]. These findings underscore the evolutionary conservation of aging pathways across diverse species.

The budding yeast *Saccharomyces cerevisiae* serves as a valuable model for studying aging, providing powerful tools for genetic manipulation in a unicellular organism with a short lifespan [5]. Aging is modeled in yeast cells by analyzing their replicative or chronological lifespans. The replicative life span (RLS) refers to the number of mitotic cell divisions that a mother cell can undergo, while the chronological life span (CLS) is defined as the number of days a non-dividing population remains viable in stationary phase. CLS is conventionally measured by following changes in colony-forming units (CFU) as a function of time in stationary phase cultures [6–9]. Using alternative high-throughput approaches, the entire yeast genome has been successfully surveyed for CLS genetic factors [10–17]. These studies in yeast highlight the roles of the TOR/Sch9 and RAS/PKA signaling pathways as conserved modulators of aging and lifespan extension by dietary restriction [12].

Despite the potential of the yeast CLS paradigm to reveal genetic players of cellular aging, a global, robust view of lifespan modulators has remained elusive. Validating genome-wide screens of CLS phenotypes has proven to be challenging, with false-positive hits ranging from 50% to over 90% when tested at the small scale by conventional CFU approaches [10–12, 14–17]. Moreover, little overlap has been documented in the lifespan modifiers identified across different genome-wide CLS screens. For instance, only nine and thirteen out of over 800 long- and short-lived mutants, respectively, are shared among three genome-wide screens [18]. Differences in media composition, genetic backgrounds, and phenotyping approaches could explain the poor consistency across results of CLS screens, especially for long-lived mutant phenotypes.

In this study, we aimed to identify genes consistently recognized as lifespan modulators in large-scale CLS screens, along with their associated features in *S. cerevisiae* and other model organisms. Using meta-analysis and functional associations within the resulting dataset, we obtained a unified, ranked catalog of genes and functions that impact CLS regardless of conditions or experimental setups. We evaluated the distribution of genes associated with RLS phenotypes and aging orthologs in animals, revealing parallels with the spectrum of CLS factors. As proof-of-concept toward a genetic-systems description of chronological longevity, we experimentally measured the lifespan-epistasis interactions of downstream CLS factors with the Tec1 transcriptional regulator of the invasive-growth signaling pathway. The ensuing lifespan-epistasis landscape provided an integrated view of the regulatory wiring underlying chronological aging in yeast.

## Results

### Genome-wide screens exhibit modest overlap in the resulting CLS phenotypes

We initially focused on ten large-scale datasets of gene-knockout strains with altered stationary-phase or postmitotic survival in *S. cerevisiae* (**Table 1**). These studies came from seven different research laboratories, all using strains of the S288C yeast deletion collection background. Large-scale surveys varied in strain ploidy, amino acid auxotrophies, culture conditions, and media composition; differences in the screening methodology were also present. A compilation of the datasets is provided in **Table S1** (see Materials and Methods). The ‘Burtner’ dataset (538 genes) was substantially smaller than the rest and was thus not considered for further analyses. CLS phenotypic data was available in at least one of the remaining nine dataset for 4779 deletion strains (**Figure S1**).

**Table 1.**
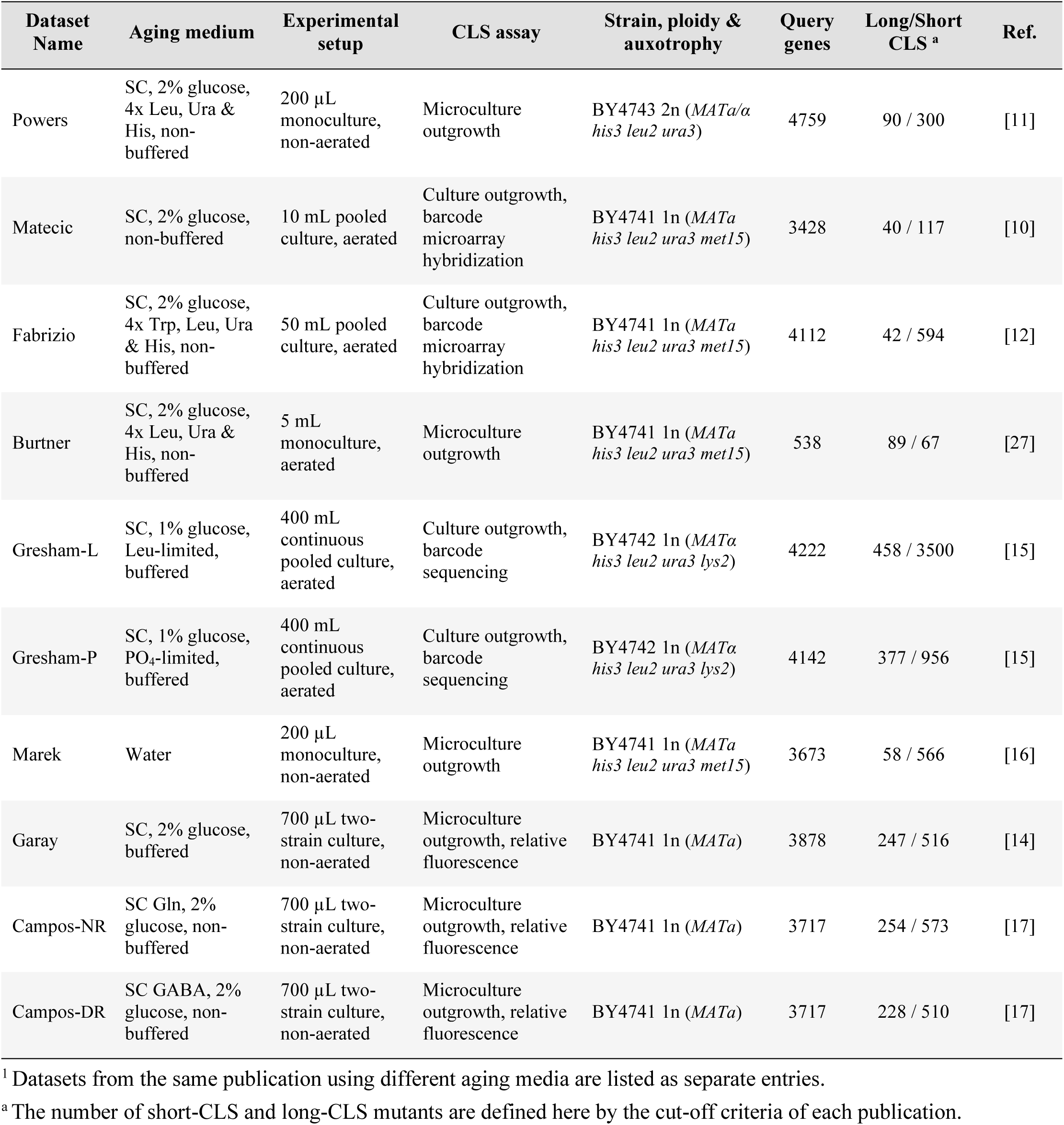
Datasets used from gene-deletion screens of CLS phenotypes in *S. cerevisiae* ^1^.

To evaluate the agreement across studies, we compared the pairwise correlation of the CLS results available in all datasets (1751 strains, 36.6% intersection). Since each study reports CLS phenotypes using different quantitative scales, we used their ranked phenotypes, from the most long-lived to the most short-lived mutant in each dataset. In agreement with previous observations using three genome-wide screens [18], modest to no rank correlation was observed between the datasets (**Figure 1**; ρ≤0.54). The highest pairwise similarities were observed among the ‘Garay’, ‘Campos-NR’, and ‘Campos-DR’ datasets (ρ=0.48 – 0.54, *p*<10^-99^), as expected given the same genetic background and experimental approach used in these studies. Likewise, the ‘Fabrizio’ and ‘Matecic’ datasets were highly correlated with each other (ρ=0.31*, p*<1.1×10^-39^), in agreement with the similar microarray barcode approach used. Given the high frequency of gene deletions with neutral or nearly neutral phenotypes, we also compared the overlap among cutoffs of long- and short-lived strains, estimated from their Jaccard similarity coefficients. This analysis confirmed that there was only a modest overlap in the scored sets of CLS phenotypes across screens (**Figure S2**). For instance, the Jaccard overlap index ranged between 0.04 and 0.29 for the most stringent cutoff for long-lived strains, while the corresponding indexes were between 0.1 and 0.38 for short-lived strains. Overall, higher overlap was observed in top ranked short-CLS in comparison to long-CLS phenotypes, as previously observed by Smith *et al*. [18]. These findings confirm that genome-wide CLS-phenotype screens exhibit only modest overlap, even when experimental conditions differ only slightly.

**Figure 1.**
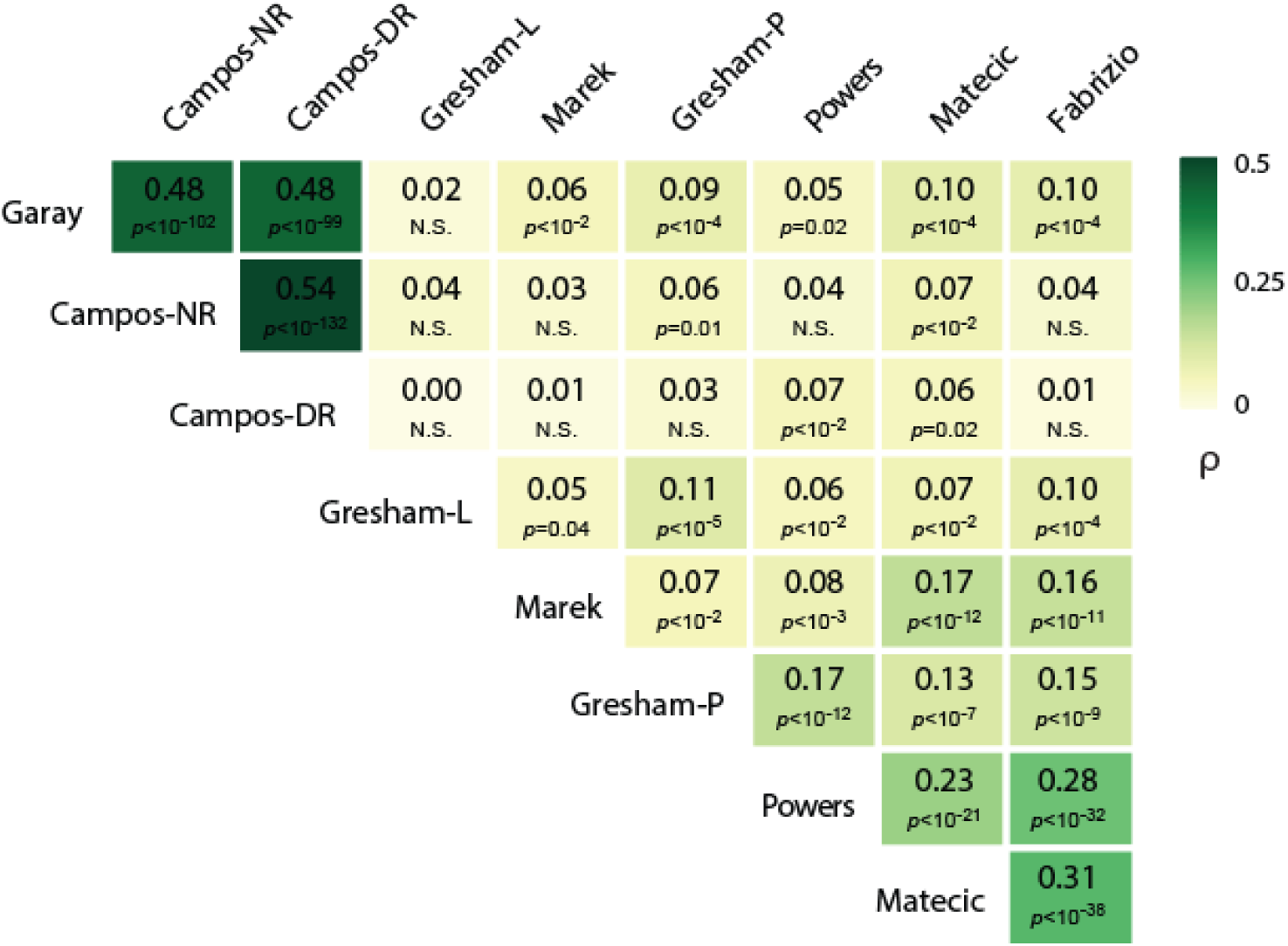
Modest correlation among genome-wide CLS assays. Spearman rank correlation matrix of the CLS datasets; mutants were ranked from highest to lowest lifespan according to each assay’s measurement and their pairwise Spearman indexes and associated *p*-values were calculated. This analysis considered only the set of genes that were shared across all datasets (n=1751). Studies are sorted in the figure by hierarchical clustering of their correlation indexes.

### A ranked catalog of genes robustly influencing chronological lifespan in yeast

We assumed that there are key mediators of chronological longevity that impact survival regardless of the conditions or experimental setup, though their relative impact may vary. To identify such a ‘core’ set of genetic factors with robust association to CLS phenotypes, we applied the PROMETHEE outranking approach [19, 20] to the CLS datasets. This method prioritizes the ranking of a set of alternatives based on a preference score (*phi-score*). These scores are obtained by pairwise comparisons of the alternatives (genes) within the criteria to be maximized (CLS phenotypes across datasets). This approach would potentially reveal, for instance, gene deletions with consistently mild CLS effects across most assays that may not have cleared the thresholds imposed by each screen. Importantly, the method is also amenable to the intrinsic differences in the quantitative units of CLS measurements and the fact that different sets of mutants were tested in the studies [21].

To establish the relative relevance of each dataset in the final priority ranking, we compiled a list of 97 gene deletions with CLS phenotypes that had been scored in smaller-scale studies using standard CFU [6] or live/dead staining approaches [7] (**Figure S3**; **Table S2**). We used this curated set of CLS phenotypes to assess the predictive performance of each of the genome-wide screens. Specifically, we calculated a receiver operating characteristic (ROC) curve for each CLS dataset, where dataset performance was evaluated by accurately classifying the lifespan phenotypes within the curated CLS phenotype set (**Figure S3**). Subsequently, we used the five datasets which showed the best predictive performance, based on their ROC curves; only one dataset per study was included for a more extensive set of conditions. Finally, by assigning proportional weights to each dataset based on how well the phenotypes recapitulated within the evaluation set (see Extended Methods in **File S1**), we obtained a final ranked list, which included all genes with available phenotypes in at least one of the five datasets used in our PROMETHEE outranking approach. We note that alternative rankings based on the different weight-allocation parameters were also analyzed, which resulted in similar rankings at the extremes of the ranks (**Figure S4**; **Table S3**).

Our meta-analysis approach resulted in a final ranked list of 4779 CLS phenotypes. The ensuing CLS longevity rank provides a genome-wide picture of genes influencing the chronological aging of yeast in a consistent manner, regardless of the experimental setup (**Figure 2A**; **Table S3**). Importantly, the assigned outranking *phi-score* of each gene reflected the weight of its phenotype relative to other genes across studies. For instance, top-ranked genes with high *phi-score*s were those with more robust evidence of increased longevity when deleted, which included *TOR1* (ranked 2.1%), *GLN3* (ranked 3.3%), and other genes involved in TOR signaling (**Figure 2A**, right panel). Genes within the lowest longevity ranks and negative *phi-score*, such as *RIM15* (ranked 99.9%) and several autophagy (*ATG*) genes, were more likely to be short-lived mutants. As expected for the postmitotic survival phenotypes herein analyzed, genes associated with the mitotic cell cycle showed a uniform distribution in the rank, with no evident enrichment at either extreme of short- or long-lived phenotypes. These general features of the ensuing ranked catalog suggest that it provides a valuable resource to interrogate the functional features robustly associated to CLS genetic factors in yeast.

**Figure 2.**
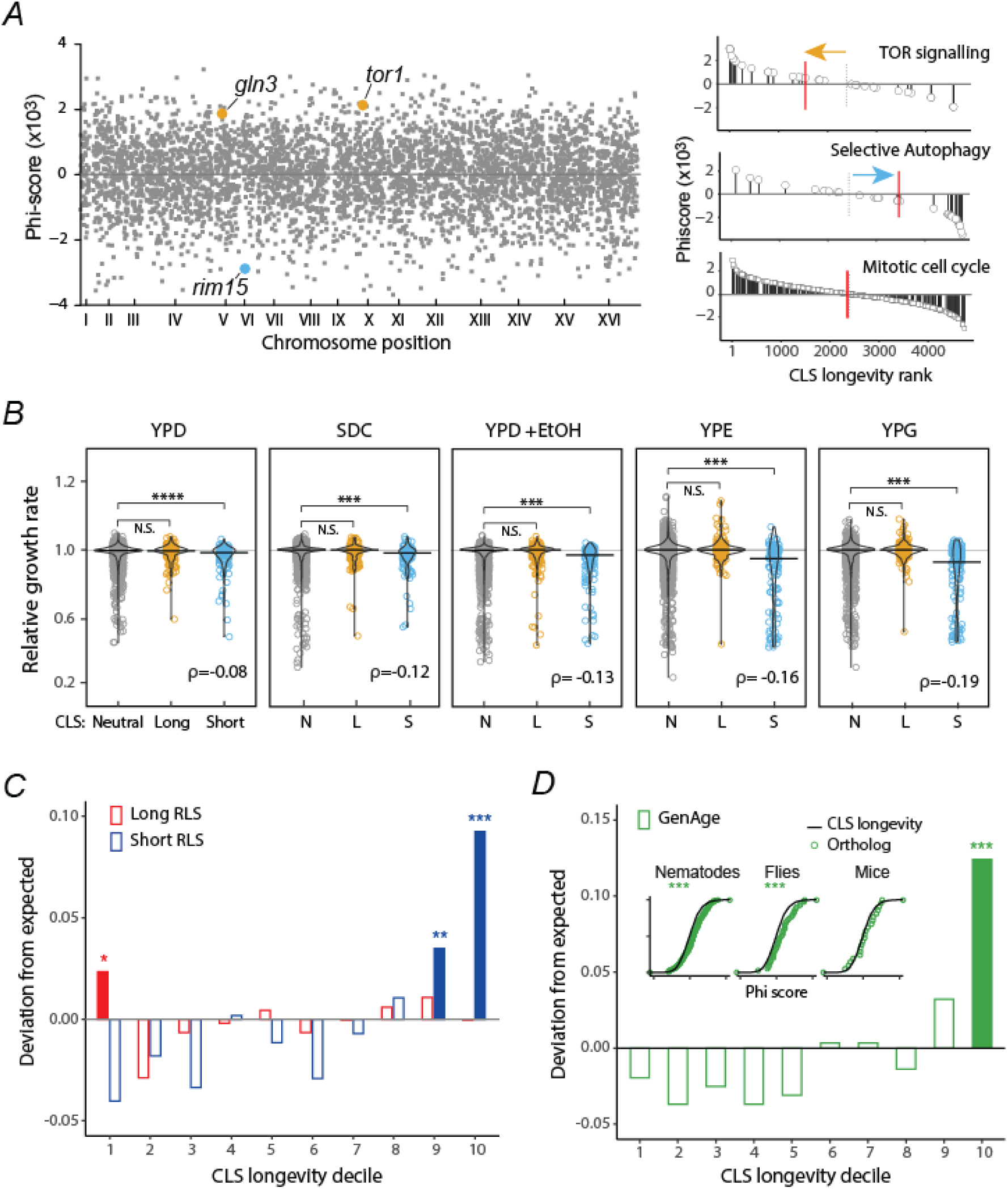
A ranked catalog of CLS phenotypes for budding yeast. **A)** *Phi-score* of genes across the yeast genome, reflecting evidence of increased (*phi*>0) or decreased (*phi*<0) longevity upon gene deletion (left). Some CLS modulators are highlighted: *TOR1* and *GLN3* (long-lived mutants) and *RIM15* (short-lived mutant). Panels on the right show the *phi-score* of genes within three functional categories; red lines are the median rank within each set; colored arrows indicate deviation from these median values. **B)** Relative growth rates of strains grouped by their inferred CLS phenotypes. Long (*n*=208) and short (*n*=122) lifespan mutants were defined as the 5% (long CLS) and 95% (short CLS) genes in the CLS longevity rank, while the rest were classified as neutral (*n*=3016) \*\*\**p*<10^-3^ Wilcoxon rank sum test. For each comparison, the Spearman correlation (ρ) is between all CLS ranks and their associated growth rates. Mutants’ growth-rate data are from [23]. Media, YPD is yeast peptone dextrose rich, SDC is synthetic complete dextrose, YPD+EtOH is 6% added ethanol, YPE is ethanol carbon source, and YPG is glycerol carbon source. **C)** Decile distribution of mutants with long RLS (red, *n*=632) or short RLS (blue, *n*=452) in the CLS longevity rank, plotted as their deviation from a null expectation of uniform distribution. Solid bars indicate deciles with a significant deviation from expected gene proportion (\*\*\**p*<0.005, \*\**p*<0.01, \**p*<0.05, hypergeometric distribution). **D)** Decile distribution in the CLS rank of *C. elegans*, *D. melanogaster*, and *M. musculus* orthologs from GenAge (*n*=134, 40, and 21, respectively), plotted as their deviation from a null expectation of uniform distribution. Solid bars indicate deciles with a higher-than-expected gene proportion (\*\*\**p*<0.005, hypergeometric distribution). Insets show the cumulative distribution of orthologs *phi-*score compared to entire dataset (\*\*\**p*<10^-3^, Wilcoxon rank sum test).

### Chronological lifespan factors are usually pleiotropic

Longevity phenotypes may be associated with the impairment of functions that are important for reproductive traits, in an antagonistic pleiotropic manner [22]. Likewise, pleiotropic functions required for reproductive fitness and long-term survival would involve gene mutations with detrimental effects on both traits. To describe the trends in the relationship between CLS and cellular fitness, we used quantitative growth-rate data obtained from yeast mutants growing under different conditions [23]. Compared to the distribution of fitness effects of mutants with no CLS phenotype, the distributions of strains with short CLS phenotypes were consistently skewed to slow-growth phenotypes (**Figure 2B**). The positive correlation between slow growth and decreased survival was higher when looking at proliferation under stress (YPD +EtOH) or respiratory conditions (YPE and YPG), compared to growth under fermentative conditions (YPD and SC) (**Figure 2B**). This indicates that stationary phase survival requires a set of functions that overlap with stress response and respiratory growth. This result suggests that the CLS phenotype in yeast is partly shaped by direct pleiotropy, but not typically by antagonistic pleiotropy. Yet, we note that many mutants with consistent CLS phenotypes showed no detectable growth defects, suggesting that a considerable number of genes impact postmitotic-survival phenotypes without playing major roles in cellular proliferation.

### Chronological lifespan factors influence the replicative lifespan of yeast

A prevailing question in yeast aging genetics is whether genes that influence CLS are also involved in RLS, and if so, which are the functions affecting both traits. To interrogate the associations of CLS genes with replicative aging, we retrieved a set of deletion strains with short-RLS and long-RLS phenotypes reported in SGD phenotype ontology [24] (**Figure 2C**). We found that short RLS mutants were more frequent in the short-CLS extreme of the rank distribution, indicating that a common set of genes and functions are required to prevent both replicative and chronological aging in yeast. In agreement with this trend, short RLS mutants were less frequent in the opposite long-CLS side of the rank distribution. Likewise, mutants with long-RLS were enriched at the top of the CLS longevity rank, reflecting a common set of genes promoting both replicative and chronological longevity. We note that long-RLS strains in the second and fourth decile were depleted, which indicates apparent opposing effects in their CLS and RLS in these cases. The main biological processes enriched in the set of shared replicative and chronological longevity factors were autophagy (*p*<10^-3^, GO over-representation test), vacuole organization (*p*=0.032), and proteolysis (*p*=0.012), while cytosolic translation typically impaired both CLS and RLS (*p*=0.027), as reflected by their shared sets of long-lived mutants. Together, these results suggest that there is a substantial set of genes contributing to mitotic and postmitotic survival in similar manners in yeast.

### Chronological lifespan factors are enriched in orthologs of aging genes in animals

Genetic analysis of aging in yeast ultimately seeks to serve as a paradigm to understand aging in other organisms, including humans. Replicative-longevity pathways in yeast have been shown to be conserved across eukaryotes, but whether conserved CLS pathways impact the lifespan of other model organisms has remained elusive [25–27]. With a catalog of consistent CLS factors in hand, we asked whether CLS factors in yeast were enriched in animal orthologs associated with aging and longevity, as compiled in the GeneAge database [28]. While no enrichment was observed in the high CLS longevity ranks, genes at the bottom of the longevity rank—short CLS mutants—had a higher fraction of orthologs associated with aging in three animal models (**Figure 2D**; *p*<10^-6^, hypergeometric distribution). We confirmed that the CLS phenotype probability distributions of the animal orthologs were different to the overall CLS longevity rank when looking independently at the orthologs from nematodes or flies, but not for mice (**Figure 2D**, insets). These results suggest that an enriched fraction of conserved genes underlying CLS have also conserved their aging-related functions from yeast to animals, underscoring the potential of the yeast chronological longevity paradigm in aging research.

### An integrated view of the downstream cellular processes of chronological lifespan

Our ranked catalog of aging factors allowed us to inquire which functions and pathways influence the CLS of yeast cells in a robust manner, independently of the experimental conditions used. To this end, we performed Gene Set Enrichment Analysis (GSEA) [29, 30] using the *phi-score* distribution of the CLS longevity rank. Enriched biological processes and cellular component ontologies were further clustered based on their shared gene content, portraying the main functions associated with CLS in yeast (**Figure 3A**; **Table S4**).

**Figure 3.**
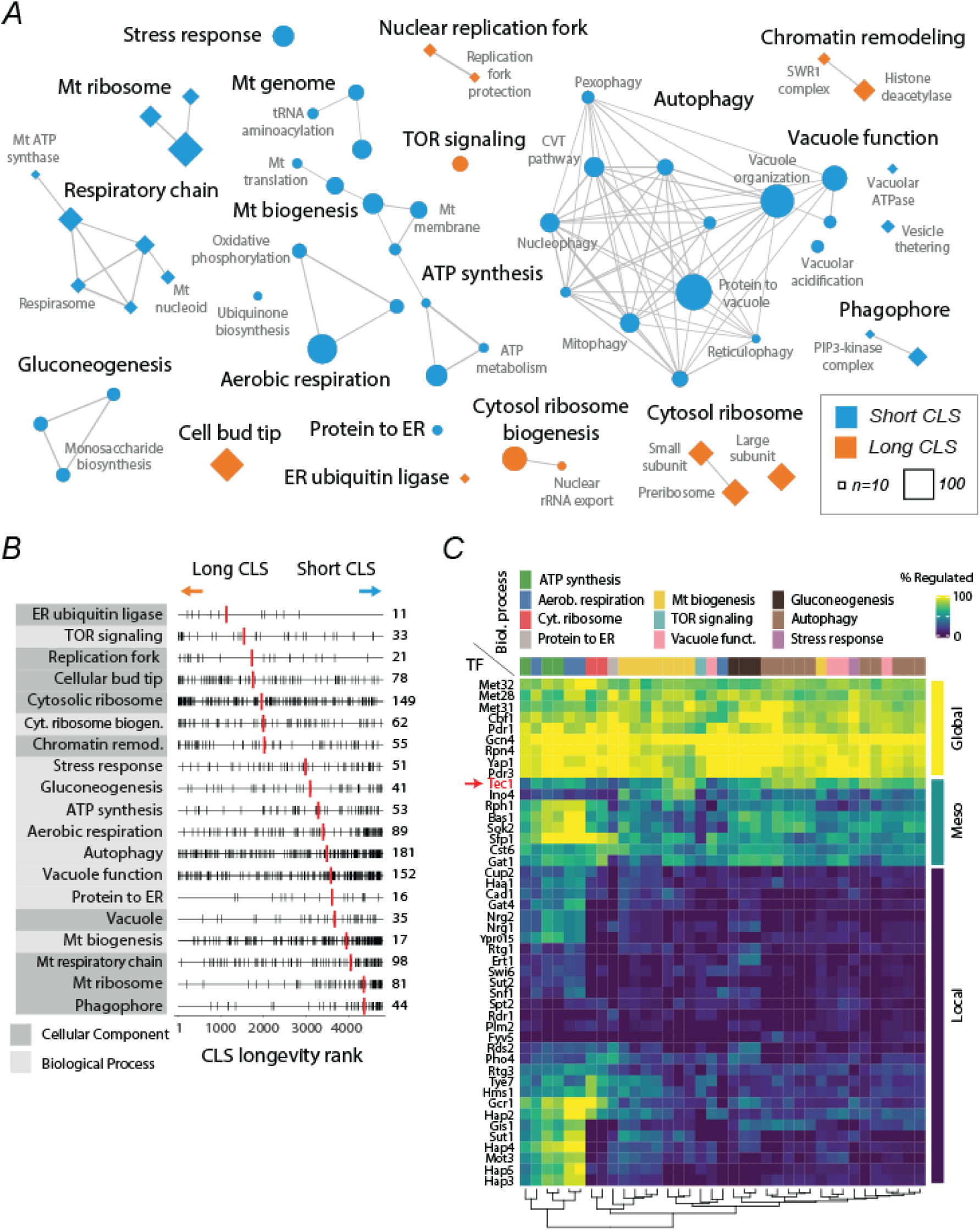
Core genetic determinants and regulators of CLS in yeast. **A)** Network representation of functional ontologies enriched at the extremes of the CLS longevity rank, according to GSEA. Each node is a Gene Othology Biological Process (circles) or Cellular Component (diamonds) term showing enrichment of long (orange) or short (blue) CLS ranks; node size indicates the number of genes within the GO term. Edges are the Jaccard indexes (*Ji*) indicating overlap in gene content between terms; only edges with *Ji*>0.2 are shown. The CLS clusters are labeled in bold according to the most common GO terms within the cluster (Table S4). **B)** The rank distribution of genes in each CLS cluster along with the median gene rank of the cluster (red lines). The number of genes in each CLS cluster is shown on the right. **C)** Heatmap showing the percentage of leading-edge genes from each Biological Process (columns) regulated by each transcription factor (rows), defined by TFRank. The matrix is sorted by the hierarchical clustering of the inferred regulatory profiles. Transcription factors were defined as ‘global’, ‘meso’, or ‘local’ regulators based on the average proportion of leading-edge genes regulated. The meso-regulator Tec1 addressed in the following section is highlighted in red.

Functional analysis showed that, in yeast, respiratory function, biogenesis of mitochondria, the ATP-synthesis machinery, autophagy and different kinds of selective autophagy, other processes involving vacuolar function, transport in the endoplasmic reticulum, response to stress, and gluconeogenesis were all consistently required for postmitotic survival, preventing premature chronological aging. At the other side of the CLS phenotype spectrum, genetic perturbations in TOR signaling, replication of DNA, protein complexes at the cell-bud tip, biogenesis of cytosolic ribosomes, and some chromatin-organization complexes usually resulted in increased chronological longevity. We observed that the resulting functional clusters associated with CLS were usually composed of genes with a wide range of phenotypes (**Figure 3B**). For instance, the cytosolic ribosome gene set was enriched at the long-CLS extreme of the spectrum but also included a considerable number of genes at the opposite end of the short-lived phenotype distribution. Likewise, autophagy and vacuole function were gene sets enriched in short-lived phenotypes, but also showing genes in the top longevity deciles. Our functional analysis of consistent CLS factors provides an integrated view of the downstream mechanisms mediating chronological aging in yeast cells.

### Transcriptional regulators of chronological lifespan factors

To provide a regulatory picture of chronological longevity, we explored which transcription factors (TFs) operate upstream of the identified CLS factors. Specifically, we used the TFRank algorithm [31] to obtain the transcriptional regulators of the leading-edge genes—those that contribute most to the observed GSEA results—within all GO terms of Biological Process CLS clusters (360 genes in total, see Methods). Forty-six TFs were the main regulators of the defined set of genes consistently associated with CLS phenotypes (**Figure 3C**; **Table S5**). At the phenotypic level, we observed that the individual deletions in this set of potential CLS regulators showed a wide range of CLS effects, with no significant changes in their median effect compared to the overall CLS longevity rank (*p*=0.90, KS test). These TFs included some known positive and negative regulators of CLS, such as Bas1 [10], Tec1 [17], Gis1 [32], and several subunits of the HAP heteromeric complex [33]. However, we note that most of this putative set of CLS regulators had not yet been described in terms of their possible contribution to chronological aging and longevity.

To obtain a broader picture of the regulatory landscape of chronological aging in yeast, we clustered the TF hits and the CLS biological processes based on the fraction of leading-edge genes regulated by each TF in each GO term (**Figure 3C**). We observed that some TFs were highly pleiotropic, connected to all or most genes within and across the CLS functional clusters, which we termed ‘global’ CLS regulators. For instance, Gcn4, Met28, Met31, Met32, and Cbf1 were global regulators of CLS, presumably by regulating their target genes with roles in amino-acid biosynthesis and sulfur metabolism. In contrast, some ‘local’ regulators were associated with specific CLS clusters, such as the Hap2, Hap3, Hap4, and Hap5 transcriptional activators involved in aerobic respiration and ATP synthesis. Importantly, at the intermediate level of the regulatory hierarchy, we observed a group of TFs linked to genes involved in apparently distinct functions (**Figure 3C**). For instance, Tec1 was associated with virtually all CLS clusters, with a higher proportion of regulated genes in some of the mitochondrial-biogenesis and autophagy clusters. These results provide a global view of the transcriptional-regulation landscape controlling chronological aging and longevity in budding yeast.

### Tec1 and mitochondria promote longevity in mutually compensatory pathways

With a catalog of consistent chronological aging factors and regulators in hand, the next challenge is to reveal how their functions are integrated with one another to drive CLS phenotype. Genetic interaction (epistasis) analysis in budding yeast has shown huge potential to describe patterns of functional association among large gene sets [34, 35]. As proof of concept, we set out to experimentally describe the lifespan-epistasis interaction landscape of the filamentous-growth pathway regulator, Tec1. Specifically, we assessed the genetic interactions of this regulator with other genes based on the measured CLS phenotypes of their single and double-gene knockouts. We decided to focus on Tec1 because of several reasons. First, in our TFRank analysis Tec1 stood out at the intermediate regulatory hierarchy, modestly associated with most CLS functional clusters, involving different functions. Moreover, the related TF Ste12—controlling filamentous growth and the response to pheromone—has been shown to be a positive regulator of chronological longevity by dietary restriction in yeast [17]. MAPKs operating upstream of these pathways are also clear determinants of CLS in yeast [36]. Despite the growing body of evidence linking filamentous-growth and pheromone-response signaling pathways to CLS, little is known about the specific downstream mechanisms of this functional crosstalk.

We first generated *de novo* gene-replacement deletions of *TEC1* or *STE12* and experimentally measured their CLS by following strain survival as a function of time in the stationary phase (**Figure 4A**). We observed that the Δ*ste12* single knockout showed decreased CLS under dietary restriction (*p*<0.05, ANOVA) while Δtec*1* showed a modest CLS effect compared to WT. While this effect was not significant in the measured survival curves, high-resolution competitive-aging screening confirmed the modest short-CLS effect of Δtec*1* (**Figure S5**). Furthermore, characterization of the Δ*tec1,*Δ*ste12* double deletion showed that the modest effects of *TEC1* were fully epistatic with *STE12*. These results suggest that the CLS effects of Tec1 regulation depend on its known association with Ste12 as a regulator of the filamentous-growth pathway. In addition, the small effects observed in the Δ*tec1* single knockout suggests the existence of compensatory pathways buffering the lack of Tec1 in terms of lifespan regulation.

**Figure 4.**
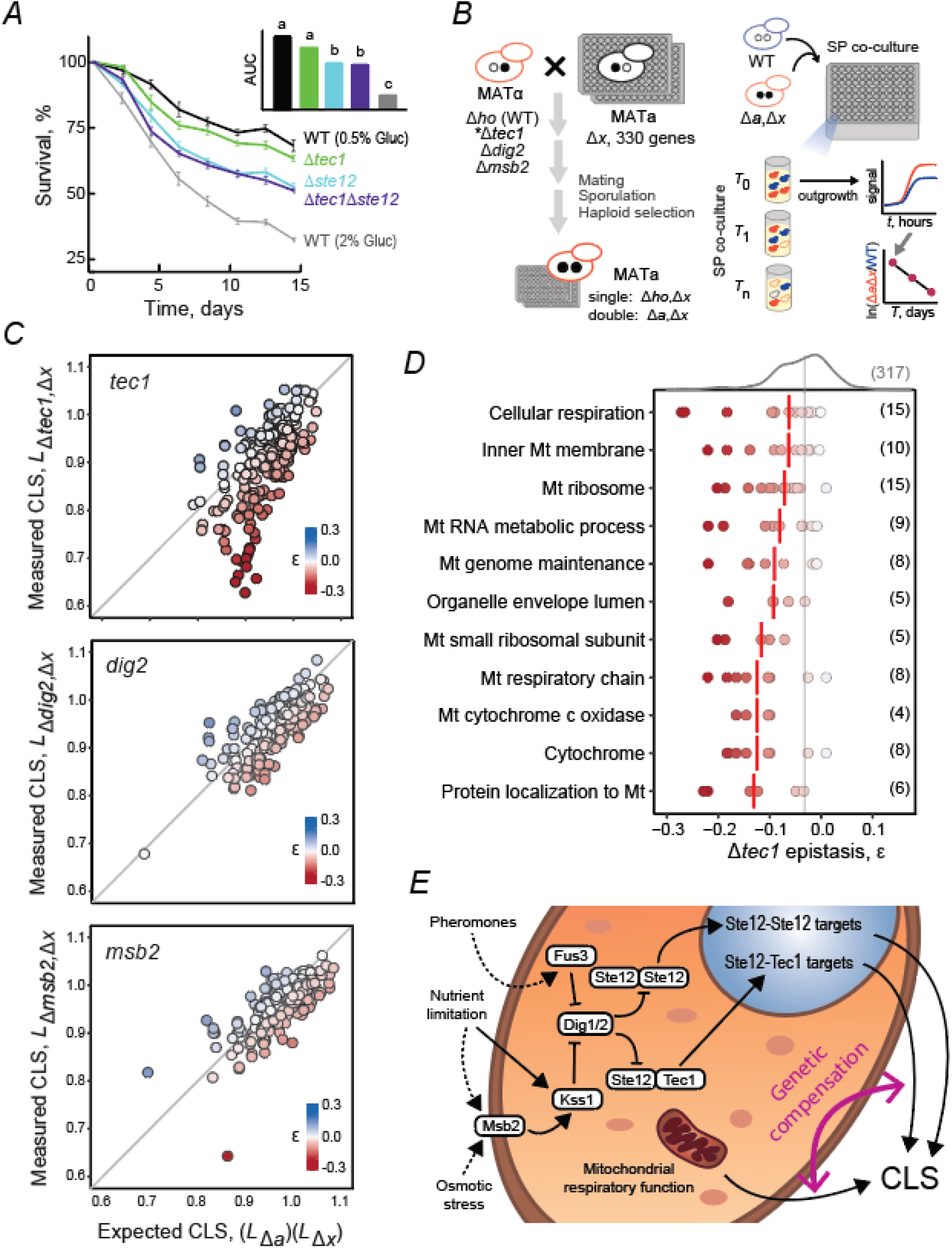
Tec1 regulation promotes longevity by compensating compromised mitochondrial function. **A)** Survival curves of WT, Δ*tec1*, Δ*ste12*, and Δ*tec1*Δ*ste12* strains aged under dietary restriction (0.5% glucose); the WT was also assessed under non-restricted conditions (2% glucose), as a reference. **B)** Schematic of the experimental competitive-aging approach used. Double knockouts were generated by mating an RFP-tagged query strain to a panel of target single deletions of representative genes of key CLS functional clusters. RFP-tagged single and double knockouts were each co-cultured in deep-well plates with a CFP-tagged reference WT strain. Stationary-phase (SP) co-cultures were analyzed by measuring both fluorescence signals in outgrowth cultures at regular time intervals as a proxy for cell viability. A relative lifespan for each mutant (*L*) was obtained by modeling the change in viable cell ratio as a function of time in the stationary phase. **C)** Lifespan-epistasis profiling of *TEC1*, *DIG2*, and *MSB2*. Scatter plots show the measured relative lifespan (*L*) of each double knockout compared to the expected *L*; data points are colored according to their epistasis (Ɛ) value. **D)** Functional analysis of *TEC1* lifespan-epistasis. Distribution of epistasis between *TEC1* and genes in the GSEA-enriched GO terms. Data series shown are all non-redundant GO terms with an epistasis spectrum that was significantly different from that of the entire target genes set (top) (*p*<0.05; Wilcoxon rank sum test). The vertical gray line is the median epistasis value of the entire target set, while each vertical red line indicates the median epistasis value of the GO term. The number of genes in each group is shown in parentheses. **E)** Functional model of chronological longevity by the filamentous-growth and pheromone-response pathways. Lifespan-epistasis analyses indicate that genes downstream of Tec1-Ste12 promote chronological longevity and compensate for compromised mitochondrial function.

To further inquire about the mechanisms underlying the effects of Tec1 as a regulator of chronological longevity and the functional nature of the pathways compensating for its genetic deletion, we carried out a synthetic-genetic array (SGA) assay [37] of *TEC1* coupled with high-resolution CLS profiling [13, 14] (**Figure 4B**). SGA targets were 330 representative genes of the key functional clusters of lifespan modulators identified in this work (**Table S6**). We note that *STE12* was not possibly used as an SGA query because the deletion strain is sterile [38], but we did include *DIG2* and *MSB2* as additional SGA queries, to compare their lifespan-epistasis profiles with that of *TEC1*. While Dig2 is a direct repressor of Tec1 and the filamentous-growth and pheromone pathways [39, 40], Msb2 elicits alternative activity of these pathways triggered by high osmolarity [41, 42]. This experimental setup resulted in the generation of a reference Δ*x* single-deletion panel and three panels of double Δ*a*,Δ*x* deletions for each of the *TEC1, DIG2*, and *MSB2* query genes (see Materials and Methods). High-resolution quantitative phenotypes were obtained using competitive-CLS profiling of all strain panels (**Figure S5**).

To screen for genetic interactions, we scored deviations of the observed and expected CLS values of each double deletion, based on a multiplicative epistasis null model. In the *TEC1* panel, we observed strong negative deviations from the expected CLS-phenotype baseline, indicating that the short-lived phenotype of Δ*tec1* was usually aggravated by the lack of specific target genes (**Figure 4C**, top). This suggests that the activities of Tec1 and several CLS genes can functionally compensate for each other’s loss. Double mutants in the *DIG2* and *MSB2* panels also showed deviation from the expected CLS phenotypes, but these were in general more symmetric and moderate (**Figure 4C**, middle and bottom).

Based on the lifespan-epistasis interactions of *TEC1*, we asked whether genes in specific CLS functional clusters were associated with this transcriptional regulator. Strikingly, a strong lifespan-epistatic interaction signal was observed for most processes related to mitochondrial function, ranging from cellular respiration and cytochrome biogenesis to maintenance of the mitochondrial genome and mitochondrial biogenesis (**Figure 4D**). This result indicates that chronological longevity by Tec1 is functionally linked to mitochondrial function, in such a way that the Tec1 targets and the aerobic respiration machinery can partially compensate for each other’s functions, which are required for CLS maintenance. It also suggests that, while Tec1 may regulate many functional clusters at the transcriptional level, its CLS-promoting effects are specifically associated with mitochondrial function. Positive genetic interactions were also observed for *TEC1* (**Table S6**); such alleviating interactions usually reflect shared pathway components. Examples of mutations that alleviated the lack of *TEC1* were Δ*ras2*, Δ*bmh1*, and Δ*gln3*, whose genes are related to cellular response to nutrients. This is consistent with the fact that the filamentous-growth MAPK pathway acting upstream of Tec1 is activated by nutrient limitation [38, 43, 44].

The lifespan epistasis profile of *DIG2* showed to be partially similar to that of *TEC1* in terms of the enriched mitochondrial functional clusters, although the magnitudes of individual interactions were smaller (**Figure S5**). This suggests that Dig2 contributes to CLS in a way like Tec1, which could be due to the fact that Dig2 is a repressor of the related pheromone-response pathway [39, 40]. Conversely, the lifespan epistasis profile of *MSB2* showed no enrichment with any of the functional clusters associated with *TEC1* or *DIG2* (**Figure S5**). This suggests that any alternative targets triggered by Msb2 are either irrelevant to CLS modulation or were missing in the panel of CLS genes. Together, results of our genetic screens indicate that Tec1 promotes chronological longevity as part of the nutrient-sensing pathway and by activating target genes that are mutually compensatory with aerobic respiration (**Figure 4E**). As proof of concept, our results show that lifespan-epistasis profiling is a straightforward way to reveal functional associations of genes and processes underlying chronological longevity.

## Discussion

The inherent simplicity of the chronological longevity paradigm, combined with the vast genetic toolbox available in yeast, would make this a straightforward model for studying the genetic underpinnings of cellular aging and longevity in a more systematic manner. However, a comprehensive understanding of the CLS phenotype has proven to be challenging. For instance, Smith *et al*. observed that only 22 CLS hits were shared among three genome-wide aging deletion screens [18]. With the larger set of studies now available [10–12, 14–17], we used a meta-analysis approach to reveal mutants associated with chronological aging in yeast. The resulting genetic catalog not only identified genes that are likely to influence CLS across various experimental setups and conditions but also offered valuable functional insights into yeast aging.

Importantly, most of the ensuing consistent downstream cellular mechanisms of chronological longevity in yeast turned out to be widely recognized hallmarks of aging [45]. Specifically, mitochondrial function, autophagy, the replication fork involved in genome stability, the protein-synthesis machinery, along with nutrient-sensing and epigenetic regulators, conformed most of the integrated functional landscape underlying CLS. The chronological aging paradigm thus offers an untapped resource to systematically inquire how these universal hallmarks of aging operating at the cellular level are integrated with one another. As proof of concept, we used SGA technology coupled to high-resolution CLS profiling to screen for lifespan-epistasis interactions of the MAPK transcriptional regulator Tec1 with genes in all the identified CLS functional clusters. We expect that applying similar methods to larger gene and functional sets will enable a comprehensive understanding of the complex genetic landscape of aging cells.

It must be noted that, even though our meta-analysis pointed to cellular processes robustly associated with aging, it still holds that the phenotypic context of chronological aging is highly variable. Our results not only confirmed a general lack of overlap in CLS phenotypes across assays [18] but also showed that even within the most relevant functional clusters there was wide variation within the CLS longevity ranking. For instance, we observed that varying only one component of the aging media, but maintaining the same genetic background and experimental setup—such as in ‘Gresham-L’ vs ‘Gresham-P’ and ‘Campos-NR’ vs ‘Campos-DR’ datasets—led to highly significant yet not fully correlated outcomes (ρ=0.11 and ρ =0.54, respectively). These extrinsic factors, along with cell-intrinsic factors such as ploidy or genetic background, have a strong impact on the CLS outcomes [46–54]. Moreover, chance variation on lifespan is also a constant phenomenon observed on laboratory assays where genotype and environment are kept constant, as evidenced in yeast, fly, and nematode models [55–57]. It is likely that the extent to which each of these contextual factors impacts CLS outcomes can only be effectively addressed on a smaller scale. Yet, as with many other areas of genomics, large-scale analyses are likely to reveal useful patterns and provide integrated knowledge, contributing to a deeper mechanistic understanding of chronological longevity, as demonstrated here.

Long-lived rather than short-lived phenotypes are arguably more informative about the underlying mechanisms of aging [58–60]. Our meta-analysis revealed cellular processes and signaling pathways whose inactivation or alteration consistently resulted in increased CLS. For instance, deletion of genes coding for ribosomal proteins or ribosome biogenesis and other processes related to cytosolic translation were linked to increased CLS. This was consistent with strong experimental evidence that ribosome regulation modifies not only yeast CLS and RLS [14, 61, 62], but also the lifespan of other model organisms. RNAi treatment directed to ribosomal genes and translational regulators increases the lifespan of *C. elegans* [63, 64], while mutation of upstream regulators of translation such as the ribosomal-protein S6 kinase and 4E-BP impacts lifespan in *D. melanogaster* [65, 66]. Deletion of the ribosomal protein s6 has also been shown to increase lifespan in mice [67]. Our results also showed that, in yeast, a major limiting factor of chronological longevity involves a widely conserved chromatin remodeling complex involved in nucleosome sliding and exchange of variant histones [68, 69]. Previously, the Swc3 subunit of the SWR1C complex was recognized as a potential pro-aging factor in a microarray-based genome-wide CLS screen [10]. Loss of several subunits of this complex mimics the effects of longevity by dietary restriction [14]. Our meta-analysis further indicates that SWR1 is a robust aging modulator, whereby its genetic inactivation results in consistently increased CLS. Yet the cellular mechanisms by which the conserved histone-exchange Swr1 complex restricts chronological longevity remain unknown.

Our experimental characterization of the lifespan-epistasis profiles of Tec1 contribute to our understanding of how MAPK signaling pathways regulate aging in yeast, which was also largely unknown. Aluru et al. revealed an unexpected role in aging of Fus3 and Kss1 involved in the pheromone and filamentation pathways, whereby deleting these MAPKs leads to increased CLS [36]. Authors speculated that downstream autophagy activity underlies this phenotype along with the observed interactions with TOR1-signaling. In a previous study, our group showed that the activity of the Ste12 transcriptional regulator of the pheromone and filamentation pathways promotes chronological longevity and that *STE12* overexpression is enough to extend CLS [17]. It was reasoned that regulation of cell-cycle arrest is the main mechanism operating downstream of this signaling pathway of increased chronological longevity [70–72]. Here, we used lifespan-epistasis profiling to directly ask which of the functional determinants of CLS explain the CLS-promoting role of Tec1, presumably in association with Ste12 and the filamentous-growth pathway. As expected, lifespan-epistasis was scored with the upstream nutrient-signaling regulators, however, no interaction was observed with autophagy, suggesting that this downstream cellular process mediates CLS independently from the Tec1 pathway. Nonetheless, we observed a strong signal of aggravating lifespan-epistasis interactions with mitochondrial function, suggesting that cellular respiration and the Tec1 pathway compensate for each other’s perturbed functions, preventing premature chronological aging. Given that Tec1 is the main transcriptional regulator of retrotransposable Ty1 elements [73–75] and that Ty activity requires intact mitochondrial function [76, 77], it is tempting to speculate that dynamics of such mobile genetic elements could explain at least part of the observed genetic interaction. Indeed, Ty1 activity promotes chronological longevity [78], although the mechanisms by which these mobile elements impact aging in yeast are unknown.

Our results also suggest that the two ways in which yeast cells age—namely mitotically and post-mitotically—share common mechanisms. Specifically, we observed an enrichment of short-RLS associated genes at the short-lived extreme of the CLS spectrum, and this was also the case for long-RLS genes at the long-lived side of CLS distribution. To the best of our knowledge, this level of genetic association between RLS and CLS had not been previously observed at the global level. Considerable efforts are being directed to gain a comprehensive dataset of RLS phenotypes [79–81], which will allow a more thorough assessment of the functional relationships of this model of mitotic aging with CLS. We also found that robust aging phenotypes were more common in yeast genes with orthologs related to aging and longevity in animal models, albeit with a weaker signal and only for short-lived effects. We note that this could be in part due to the observed direct pleiotropy nature of many CLS genes, since genes with a high contribution to fitness tend to be conserved across organisms [82].

Our study highlights the main mechanisms of chronological aging in yeast, which shares many of the widely recognized universal underpinnings of aging. Moreover, the genetic relationships between RLS and CLS, along with the evolutionary and functional conservation of some aging genes from yeast to animals, underscore the potential use of the amenable yeast CLS model in aging research. By fully exploiting the vast genetic toolbox in yeast, the chronological longevity paradigm will provide further insights into the genetic logic of aging and the genetic wiring of aging cells.

## Materials and Methods

### Strains and media

Single *de novo* knockout mutants were generated by PCR-based direct gene replacement using natMX4 cassette from the pAG25 plasmid on the YEG01-RFP and YEG01-CFP parental strains (*MAT*α *PDC1-XFP-CaURA3MX4 can1*Δ:*STE2pr-SpHIS5 lyp1*Δ *his3*Δ1 *ura3*Δ0 *LEU2*) by direct gene replacement with natMX4 cassette from pAG25 [14]. For gene interaction analysis, double mutants were generated using the Synthetic Genetic Array methodology [37]. Briefly, the RFP-tagged *de novo* query strains were mated to a target array of selected mutants of the yeast deletion collection (*MAT*a BY4741) [83]. Diploids were sporulated and subjected to three rounds of haploid selection (*HIS^+^* for *MATa* mating type, *URA^+^* for the fluorescence marker, and *G418^+^* and NAT*^+^* for knockout selection).

Synthetic Complete (SC) aging medium used for non-restricted conditions contained 6.7 g/L yeast nitrogen base (YNB) without amino acids (Difco 291940), 20 g/L glucose, and 2 g/L amino acid supplement mix (Yeast Synthetic Drop-out Medium Supplements without uracil, Sigma Y1501, plus 0.076 g/L Uracil). Dietary restricted media (DR) was SC with 10 g/L glucose (0.5% glucose). Low fluorescence medium (YNB-lf) for outgrowth cultures was prepared as described [84].

### Competitive-aging CLS assay

High throughput fluorescence-based CLS profiling of single and double deletion panels was based on Avelar-Rivas *et al.* (2020) [13]. In brief, saturated cultures of the selected strains were pinned to 96-well plates with 150 μl of fresh SC medium. These plates were left to saturate for 48 h at 30°C without shaking. Saturated RFP-labeled mutants and CFP-labeled WT cultures were mixed in 2:1 RFP:CFP ratio; reference wells with WT_RFP_ or WT_CFP_ strains only were included, for background fluorescence measurements. Mixed cultures were pinned to 96-well deep wells containing 700 μl of SC medium and grown to saturation at 30°C and 70% relative humidity, without shaking. After four days and at daily intervals afterwards, cultures were resuspended by shaking and 5 μl aliquots were inoculated with an automated robotic pipetting arm into 150 μl of fresh YNB-lf medium. In these outgrowth cultures, OD600 and fluorescence (RFP and CFP) were measured every 2 h until saturation, with a Tecan Infinite M1000 plate reader. Sampling was repeated for six days in total. Optimal gain values were 164 and 204 for CFP and RFP signal, respectively; these were obtained from a previous outgrowth at late exponential growth, when the signal was at its maximum.

### Data analysis for competition-based CLS screening

Death rates for the fluorescence-based assay were calculated following Avelar-Rivas *et al.* 2020 [13]. The RFP/CFP ratio was used to estimate the number of cells tagged with each fluorescence protein. Background fluorescence was subtracted for all measurements. For each sampling day (*T_i_*, in days) the outgrowth cultures measurements (*t_j,_* in hours) were compiled into a signal ratio *ln(RFP/CFP)T_i,tj_* for each sample (w) and fitted to the linear model *A_w_* + *S_w_*⋅*T_i_* + *G_w_*⋅*t_j_* + *C_Ti_*,*t_j_*., where *A* was the viable cell ratio at the start of the experiment, *G* was the growth rate difference of both strains, and *S* was the survival of the mutant relative to the WT reference. An additional term, *C_Ti_, t_j_*, was included to model the systematic variation of each plate at each stationary-phase sampling point *T_i_t_j_*. *S*=0 and *G*=0 was assumed for all WT_RFP_/WT_CFP_ competitions. The resulting set of equations was solved for *Sw* by multiple linear regression. Relative survival (*Sw*) was transformed to relative lifespan values (*Lw*), where *Lw*= 1+*Sw*.

### Definition of epistasis, genetic interaction profiling, and functional analysis

Epistasis was calculated following a multiplicative neutral expectation model. For each deletion gene pair *x*Δ and *y*Δ, epistasis was defined as ε=*Lxy*–(*Lx*\**Ly*), where *Lx*, *Ly* and *Lxy* were the relative survival of the *x*Δ and yΔ single mutants and the *x*Δ*y*Δ, double mutant, respectively. For functional analysis, Epistasis values were averaged when replicates were available for a particular competition. Genes were grouped according to their key CLS modulator assignment, as shown in **Table S4**, then tested for significance.

A Wilcoxon rank sum test was used to determine significant interactions. Distribution of key CLS modulator epistasis was compared against the distribution of all genes tested. A key CLS modulator group was determined to have a significant interaction with the query gene using a *p*<0.05 cutoff.

### CLS characterization by outgrowth kinetics

Small-scale CLS assays were based on the outgrowth kinetics method described by Murakami *et al* (2009) [8]. In brief, selected strains were grown in aerated tubes individually in either NR or DR media for 48 h at 30°C with 200 rpm, then transferred to 96-well microtiter plates for array setup. These plates were replicated by pinning onto 96 deep-well plates containing 700 μl of NR or DR media. Plates were then incubated at 30°C and 70% relative humidity without shaking. After 4 days of growth and every 2 to 3 days for 21-28 days, 10 μl samples were taken from the culture with an automated robotic pipetting arm (Tecan Freedom EVO200) and inoculated in 96-well plates with 150 μl of fresh YNB-lf medium. OD600 was measured using a plate reader (Tecan M1000) every 1.5 h until saturation. The first outgrowth-kinetics curve was considered as the first time point (To). At each sampling time point, the time shift to reach mid-exponential phase (OD600=0.3) was extracted (Tn). Experiments were performed in at least three independent replicates and survival was calculated from these data, as described in [8].

### Chronological lifespan datasets

Genome-wide chronological lifespan assay data was retrieved for each publication mentioned in **Table 1** [10–12, 14–17, 27]; the experimental setup of each dataset is also shown. In some publications, more than one medium was used for measuring the lifespan of a given knockout mutation set. In such cases, each condition was assigned as a different dataset and was analyzed separately. To integrate all high-throughput experiments into a summarized table, for every dataset a single representative lifespan value of each knockout mutant was taken. Details of data transformation for each dataset are available as Extended Methods (**File S1**). This process resulted in ten different datasets which can be found in **Table S1**.

### Dataset evaluation and AUC-ROC score

A curated knockout strains list with a known confirmed phenotype in CLS was generated to be used as reference to evaluate data reproducibility (**Table S2**). An initial list was obtained from the Yeast Phenotype Ontology at the Saccharomyces Genome Database (https://yeastgenome.org/ontology/phenotype/ypo) [24]. The above-mentioned list was generated by filtering the phenotype ontology for “Chronological lifespan increased” or “Chronological lifespan decreased” annotations, plus the “classical genetics” annotation. Further manual inspection of the records was done to select those mutants whose phenotype was validated by using the colony forming unit (CFU) method or live-dead staining assay. Mutants were retrieved regardless of reported culture media and conditions. Mutants were eliminated from the list if they presented opposite phenotypes, long-lived and short-lived, even if those reports were made on different culture media. To assess the CLS datasets replicability, each dataset was evaluated on their performance to segregate mutants of known phenotypes in two groups: long-lived and short-lived. With the curated mutant list, a set of true labels were generated for a binary classifier system. While testing each dataset, curated mutants were excluded when the small-scale phenotypes were validated within the same study, to avoid performance overestimation due to common source bias (**Table S2**). A Receiver Operating Characteristic curve (ROC) was used to evaluate the performance of each CLS dataset, using the *ROCR* package in *R* [85]. Further details on the ROC evaluation are provided as Extended Methods (**File S1**). Datasets were ranked according to their area under the ROC curve and the top five datasets were used for PROMETHEE analysis.

### PROMETHEE method for multivariate analysis

To generate a ranking of which gene deletions have been consistently reported as long-lived, the PROMETHEE II method of multiple-criteria decision analysis was used [19, 20], which was implemented in the *RMCriteria* package in *R* (version 4.3.0) was used [86]. The CLS datasets were set as criteria for the method. Five datasets were used as ranking criteria: ‘Matecic’, ‘Marek’, ‘Campos-DR’, ‘Garay’, and ‘Gresham-L’. All genes with available data in at least one dataset were used as alternatives. For PROMETHEE II analysis settings, a usual preference function was applied to maximize all criteria. Weight allocation for final ranking was proportional to area under ROC curve. (w_1_=0.233, w_2_=0.197, w_3_=0.195, w_4_=0.194, w_5_=0.181). Net preference flow after PROMETHEE II analysis was used to generate the ordinal ranking and reported as *phi-score*s. Further details on weight allocation parameters and alternative rankings are provided as Extended Methods (**File S1**).

### Other phenotypic datasets

Growth rates from single knockout mutants under a variety of culture conditions were retrieved from Qian et al 2012 [23]. RLS data List of genes with an effect over replicative lifespan was retrieved from the Yeast Phenotype Ontology at the Saccharomyces Genome Database [24]. The records labeled under “increased replicative lifespan” and “decreased replicative lifespan” were retrieved and filtered to select knockout mutations and microdissection RLS methods. Genes associated to lifespan modulation in mouse (*Mus musculus*), fruit fly (*Drosophila melanogaster*), and nematode (*Caenorhabditis elegans*) were retrieved from the GenAge database (https://genomics.senescence.info/genes/models.html) [28]. Genes were included regardless of their short or long lifespan effects or the genetic perturbation used for phenotyping. Yeast orthologs of these genes were also retrieved from GenAge.

### Gene Set Enrichment Analysis

Functional enrichment analysis of the ranked list of robust lifespan factors was done using the Gene Set Enrichment Analysis (GSEA) method implemented on *clusterProfiler* package in *R* [29, 30]. *Phi-scores* derived from PROMETHEE analysis were used as input for gene list ranking. Biological Process (BP) and Cellular Component (CC) GO terms were assessed as gene sets for enrichment. GO term annotations were retrieved from org.Sc.sgd.db package (v.3.18) [87]. Only gene sets with 10-100 genes were considered for GSEA. GO terms with FDR-adjusted *p*> 0.05 were regarded as enriched. The Jaccard index was calculated for comparing the gene overlap among the enriched sets. Hierarchical clustering of the resulting Jaccard indexes was done to identify redundant or similar GO terms. Assignation of GO terms to biological clusters was based on this clustering, using arbitrary names. Full and revisited list of enriched gene sets along with functional clusters are provided in **Table S4**.

### TFRank analysis

Prediction of the main transcriptional factors regulating CLS consistent biological processes was made using the TFRank approach (http://www.yeastract.com/formrankbytf.php) [31]. Leading Edge genes of enriched GO terms of GSEA in the BP category were used as input (**Table S4**). Settings were set to check for all TFs available in YEASTRACT and filter documented regulations by DNA binding or expression evidence. No filtering by environmental conditions was added. TFs were considered of relevance if *p*<0.01. Significant TFs, their targets within leading edge genes, and percentage of genes regulated are provided in **Table S5**.

## Acknowledgements

We thank Mayra Flores-Barraza and Ariann E. Mendoza for technical support and Diana Ascencio and J. Abraham Avelar-Rivas for critical reading of the manuscript. This work was funded by the Secretaría de Ciencia, Humanidades, Tecnología e Innovación de México (Secihti), Grants CB2015/254365, CF-2009-G-103000, and CF-2023-I-1545. E.C-B. was funded by doctoral fellowship CVU 711011 from Secihti. A.D. was funded by a sabbatical fellowship CVU 26244 from Secihti.

## Contributions

E.C-B. and A.D. conceptualized and designed the study. E.C-B. and S.C. performed experiments. E.C-B. performed data analysis. S.F. and A.D. oversaw experiments. C.A-G. and A.D. oversaw data analysis. S.F. and A.D. acquired funding. E.C-B. and A.D. wrote the original draft. All authors read and critically revised the manuscript.

## Supplementary Material

**File S1.** Extended Methods.

**Table S1**. Datasets used from gene-deletion screens of CLS phenotypes in *S. cerevisiae*.

**Table S2**. Curated CLS phenotype set used for evaluating the chronological lifespan datasets on AUC-ROC curves.

**Table S3**. CLS longevity rank obtained from metanalysis of CLS high-throughput assays.

**Table S4.** GSEA results of GO terms enriched in extremes of CLS longevity rank.

**Table S5.** Transcriptional factors regulating leading edge genes of CLS functional clusters.

**Table S6.** Relative CLS and epistasis of genetic interaction assays of *TEC1*, *DIG2*, and *MSB2*.

---

**Supplementary Figure S1.**
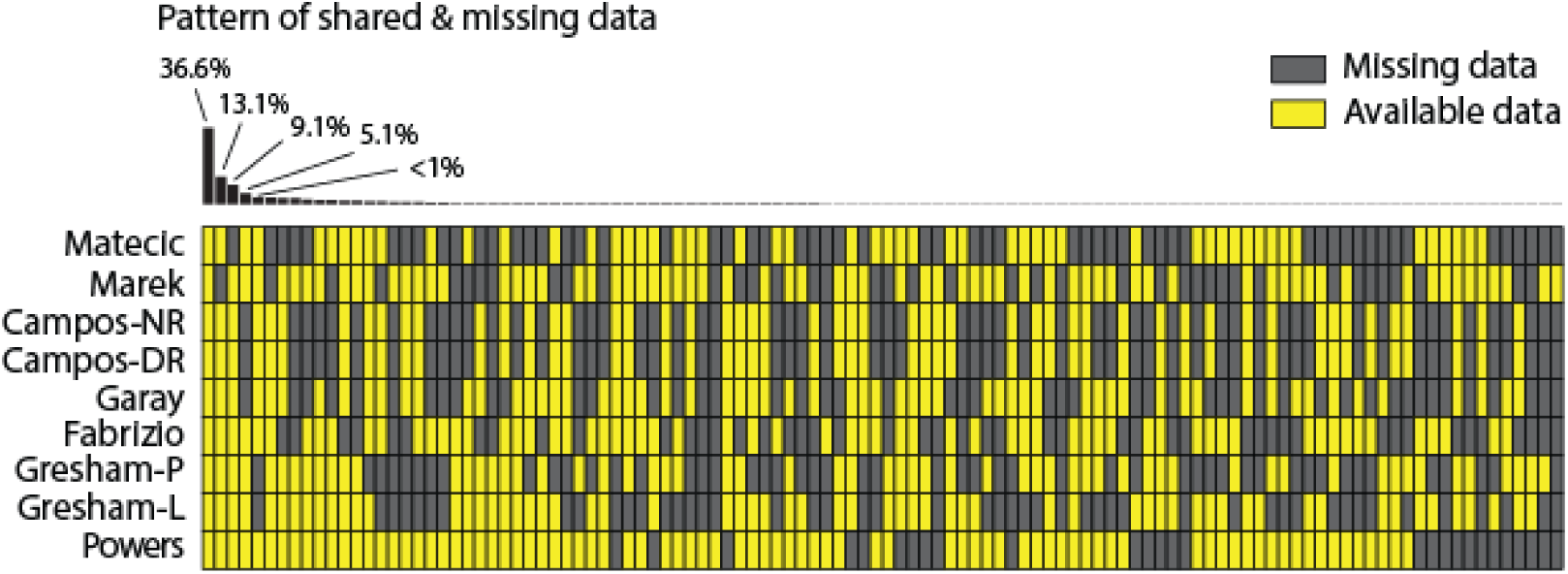
Comparison of data available on the high-throughput CLS datasets. Data availability patterns of knockout mutants are shown for all datasets, except ‘Burtner’ due to the large fraction of missing data. The most comprehensive dataset was ‘Powers’ (1.7% missing genes), while the least complete were ‘Gresham-P’ with 13.1% and ‘Matecic’ with 9.1% missing genes. Gene availability patterns are labeled with their corresponding percentage; patterns with no percentage shown account together for ∼13% of occurrence patterns, but each with less than 1% occurrence.

**Supplementary Figure S2.**
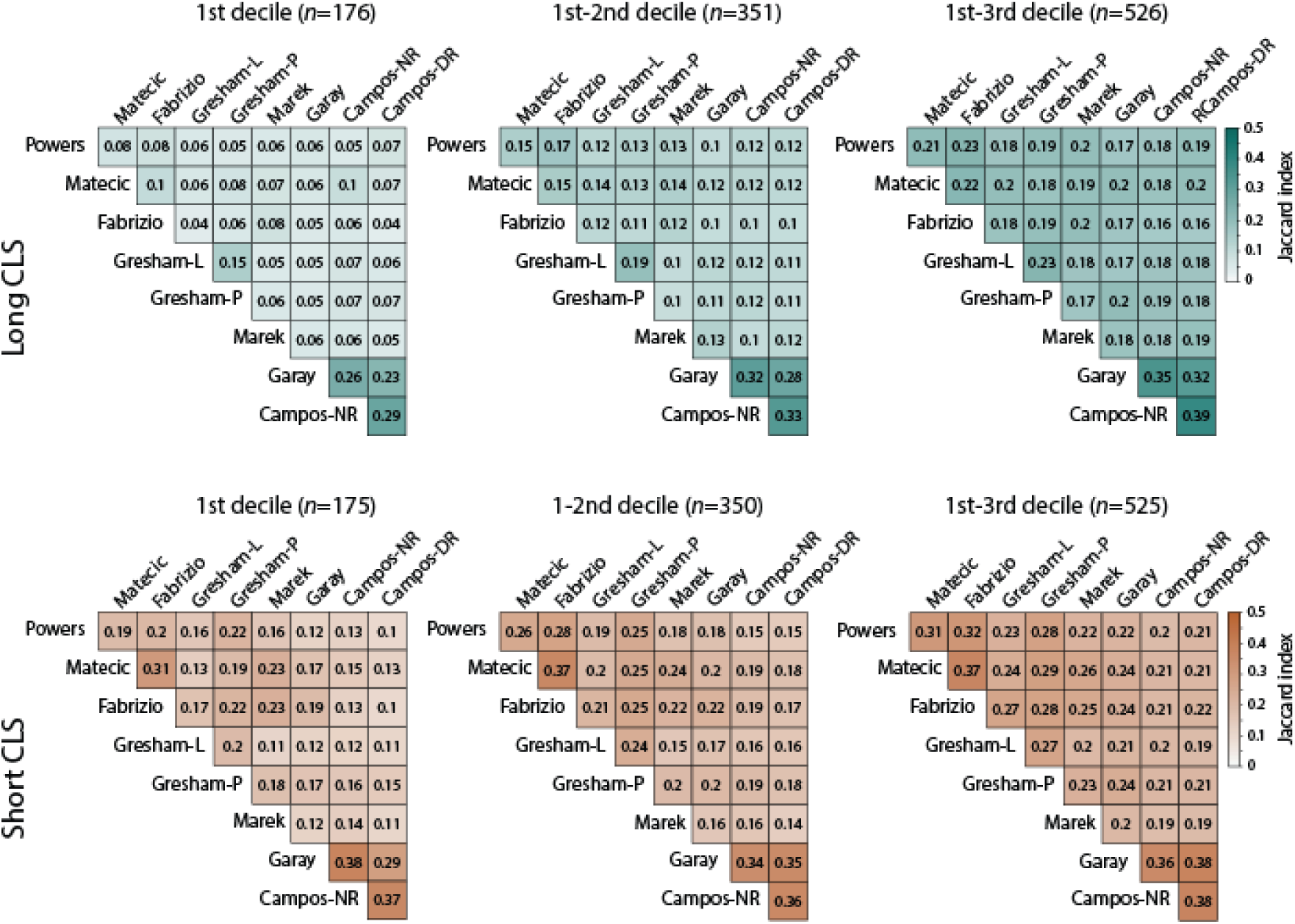
Overlap similarity among CLS datasets. Each ranked list of mutants, from highest to lowest lifespan according to each assay measurements, was divided into deciles. Only the shared gene set among all datasets is considered in this comparison (n=1751). The first three deciles were assigned as long-lived mutants, while the last three were defined as short-lived. Overlap among dataset deciles was estimated with the Jaccard similarity index. The number of genes in each decile comparison is shown.

**Supplementary Figure S3.**
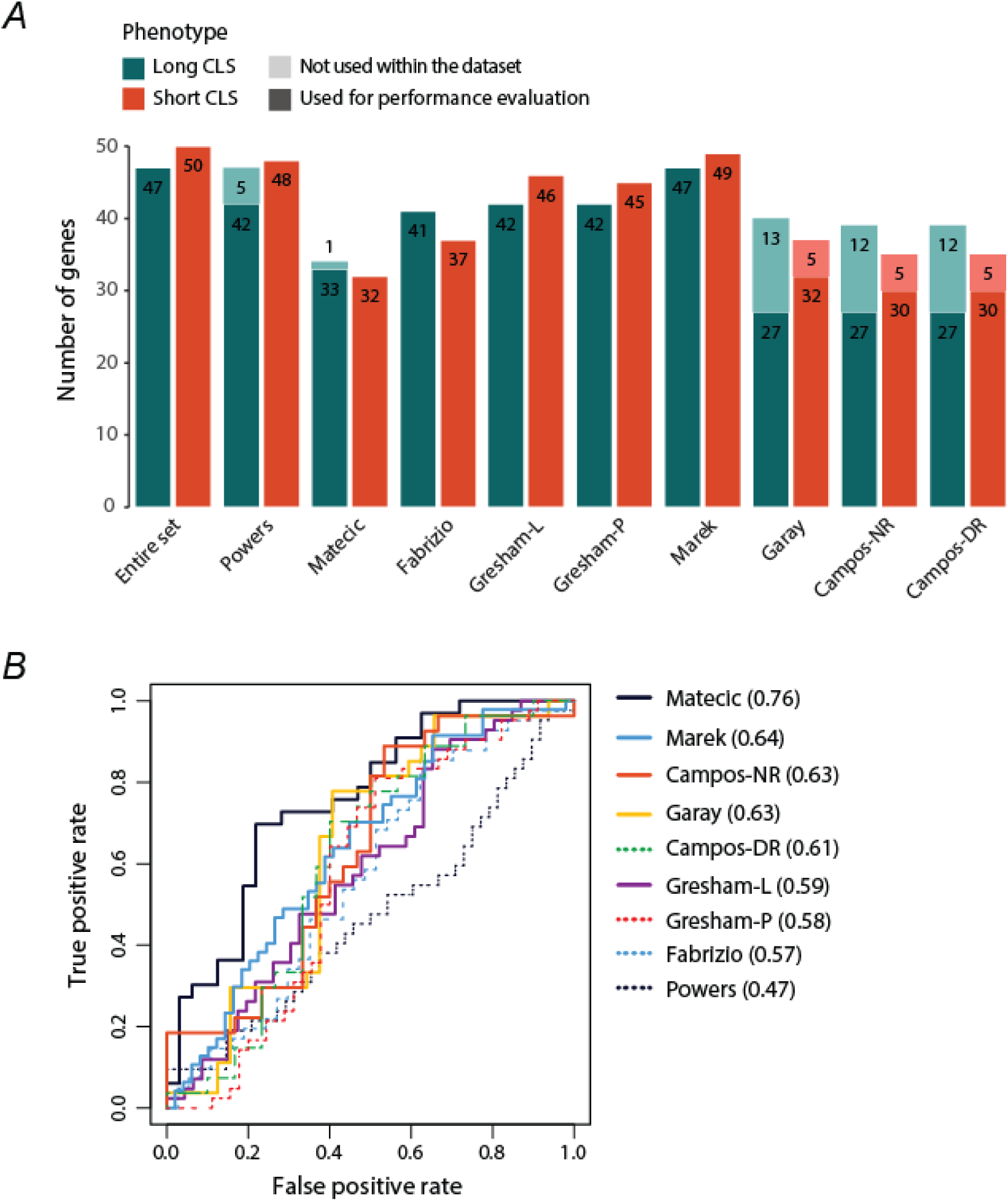
Evaluation of CLS datasets on AUC-ROC curves. **A)** Availability of the curated CLS phenotype set used to evaluate the datasets. Mutants are separated into long- and short-lived phenotype. We note that the curated CLS phenotype set includes mutants that were validated at the low scale in some of the large-scale screens evaluated (light background in bars). The final number of mutants used for ROC evaluation after common source removal is shown in solid background. **B)** ROC curves of high-throughput CLS datasets against the curated CLS phenotype set. The area under curve (AUC) is shown beside each dataset. Datasets selected for the final ranked list of robust CLS factors are shown with solid lines.

**Supplementary Figure S4.**
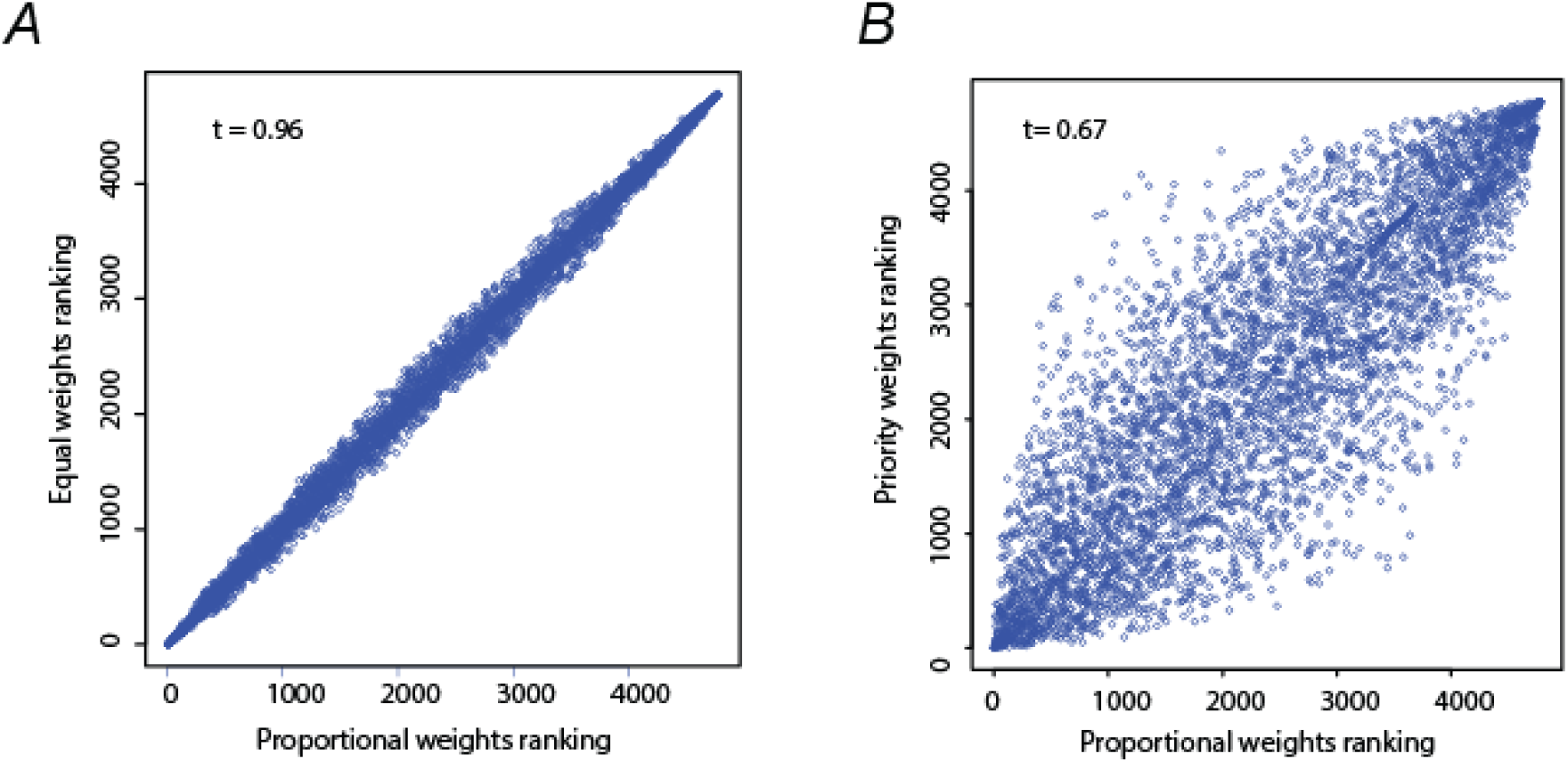
Comparison of alternative rankings obtained using different weight allocations for the CLS datasets on the PROMETHEE II method. Ranking with original weight allocation is labeled here as Proportional. We generated two alternative rankings with new weight distributions, labelled Equal and Priority. Weight allocations for each ranking can be found in Extended Methods (File S1). Kendall tau correlation is shown in each comparison between **A)** proportional vs equal and **B)** proportional vs priority weights rankings.

**Supplementary Figure S5.**
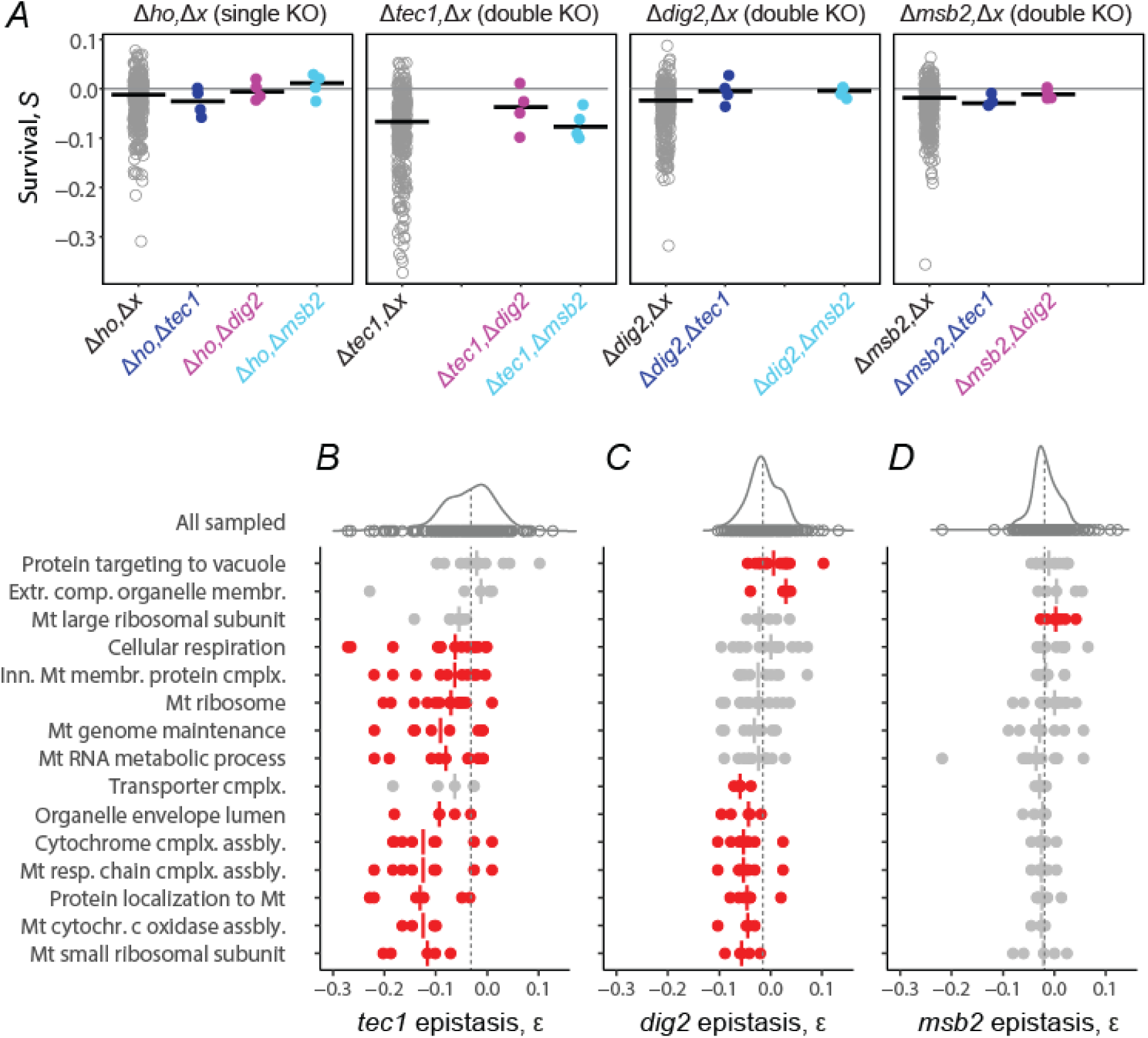
CLS Genetic interaction assay phenotypes and interaction profiles of *TEC1*, *DIG2*, and *MSB2* genetic interaction assay. **A)** Survival coefficients (*S*) of all mutants phenotyped for genetic interaction assays. Mutants are grouped based on their genotype on single (*Δho,Δx*), Tec1 (*Δtec1,Δx*), Dig2 (*Δdig2,Δx*) and Msb2 (*Δmsb2,Δx*) double mutants. CLS of query genes single and double mutants is shown apart. The median *S* is shown for each category. Genetic interaction profiles of **B)** *TEC1*, **C)** *DIG2*, and **D)** *MSB2*. Epistasis distribution of GSEA-enriched GO terms with significant bias in at least one genetic interaction assay are shown, but significant GO terms in a particular interaction assay are colored red (*p*<0.05, Wilcoxon rank sum test). Median epistasis for each GO term is indicated with vertical lines. Epistasis distribution of all genes sampled in each interaction assay is also shown (top).

